## Supplemental Figures for "Single-cell multiomic human brain atlas reveals regulatory drivers of cortical regionality"

**The file includes:**

Supplementary Figs. 1-14

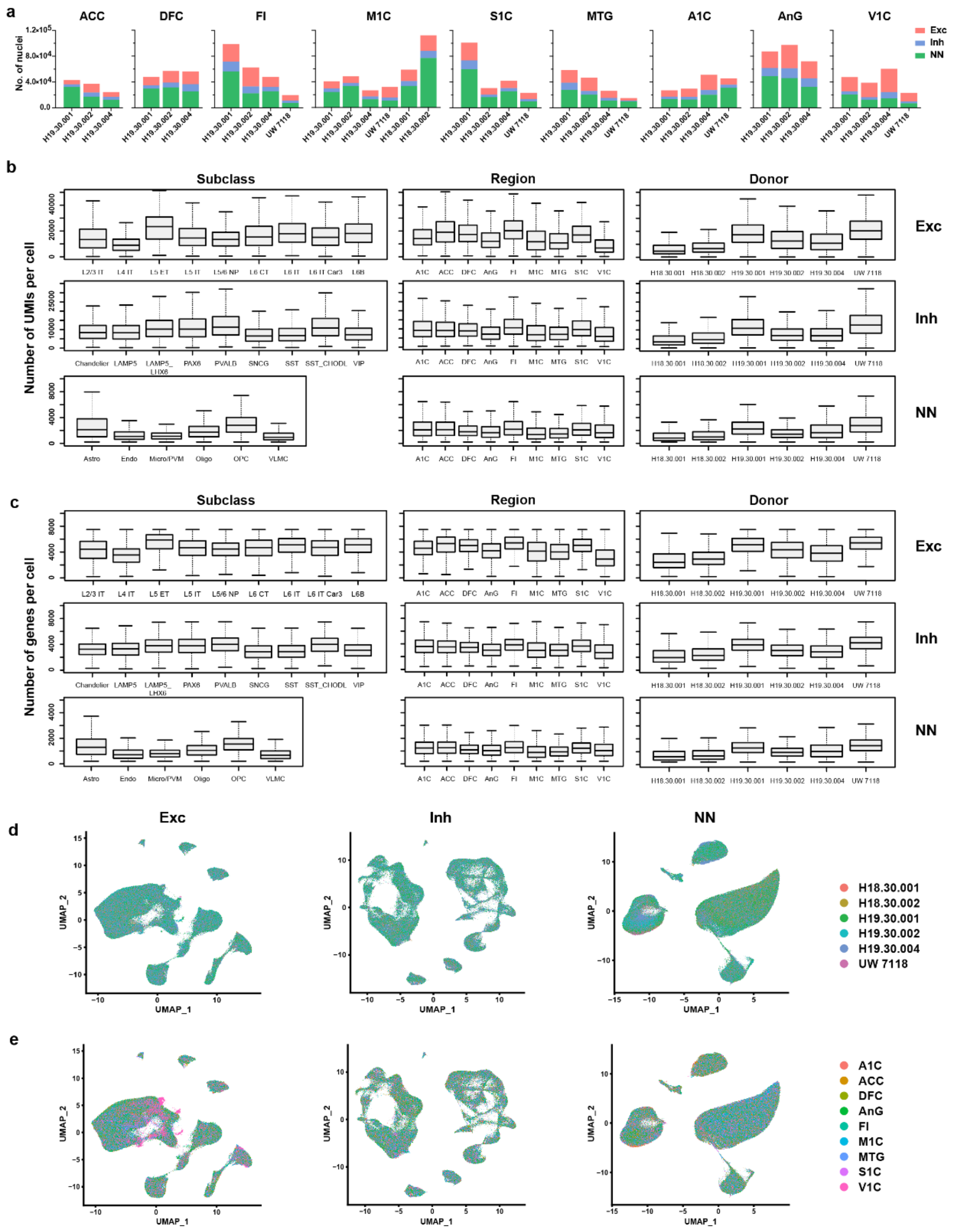


**Supplementary Figure 1.**

**Quality control metrics for single-nucleus RNA sequencing data.** (**a**) Bar plots showing the number of nuclei that passed QC for each region and each donor. (**b**) and (**c**) Box plots showing the UMIs per cell (**c**) and genes per cell (**c**) in each subclass, region or donor. (**d**) and (**e**) UMAP embedding and clustering analysis from RNA data of three major classes, including excitatory neurons (left, Exc), inhibitory neurons (middle, Inh) and non-neurons (right, NN). Individual nuclei are colored and labeled by donors (**d**) or regions (**e**).

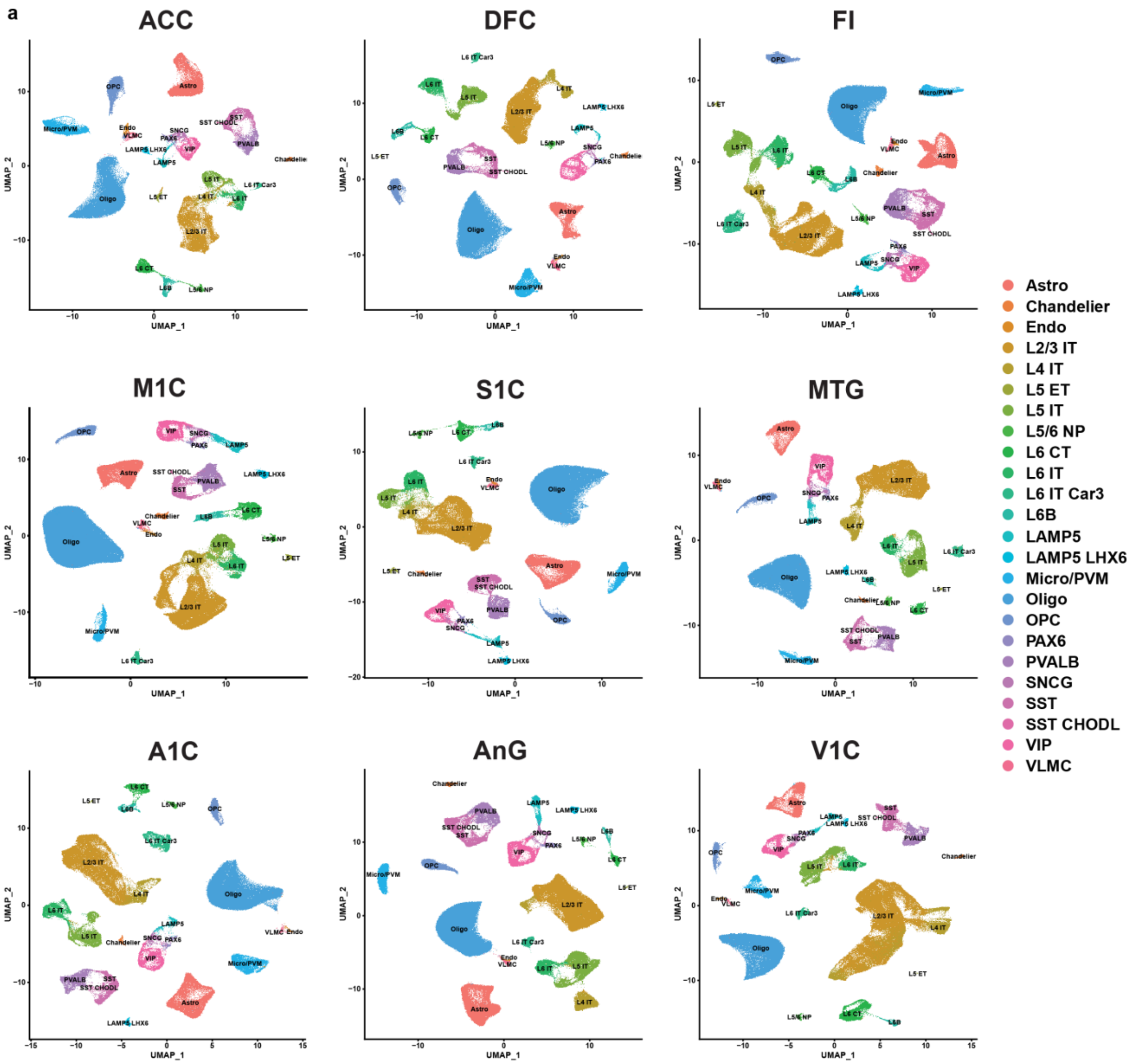


**Supplementary Figure 2. RNA-based UMAP analysis by region.** (**a**) UMAPs based on variable gene expression and colored by cell subclasses for each profiled region.

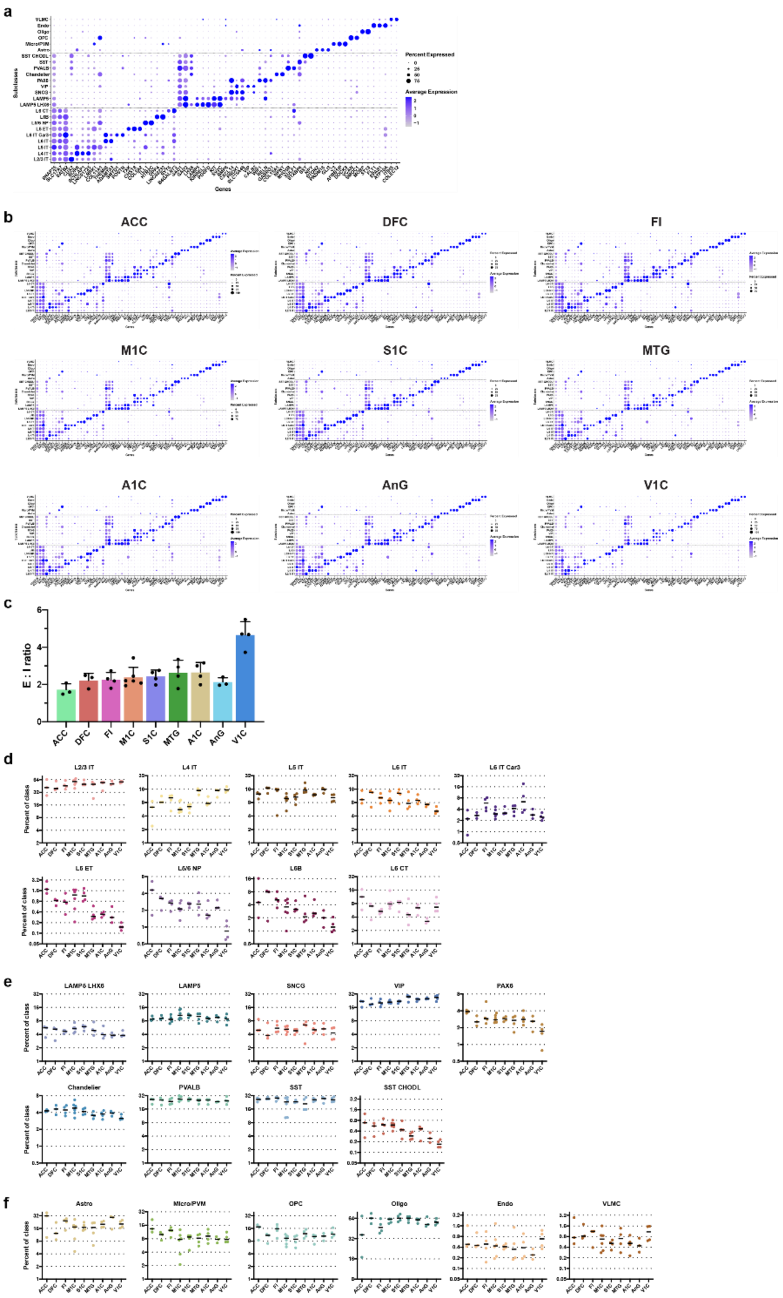


**Supplementary Figure 3.**

**Cell subclass composition and maker genes of adult human cortex.** (**a**) Percent expressed cells and expression level of representative marker genes. Detailed marker information is provided in table S3 and S4. (**b**) Percent expressed cells and expression level of representative marker genes after analysis for each region individually. (**c**) Bar plots showing the ratio of excitatory neurons to inhibitory neurons (E:I ratio) in each region. Data are mean ± standard deviation across donors. Each dot represents a sampled brain. **(d-f)** Subclass proportions for each donor in each major class, including excitatory neurons (**d**), inhibitory neurons (**e**) and non-neurons (**f**).

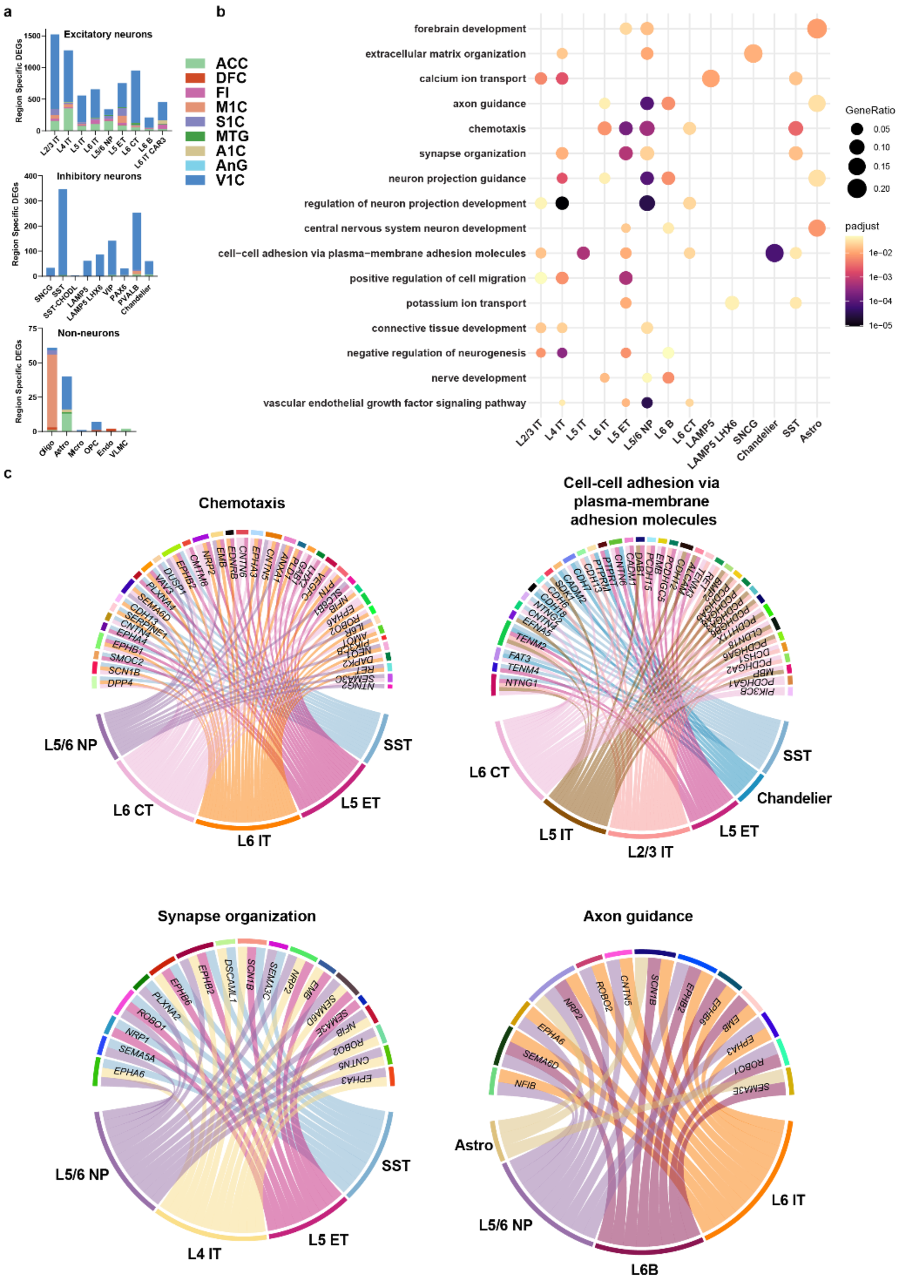


**Supplementary Figure 4. Gene Set Enrichment Analysis of individual subclass regional specific differentially expressed genes.** (**a**) Total counts of region-specific differentially expressed genes (DEGs) calculated with edgeR as genes differentially expressed in one region as compared to all others combined. Variable DEGs calculated from all cells that passed filtering. Missing subclasses had zero region-specific DEGs. (**b**) Dot plot showing both gene ratio and adjusted p-value for top enriched terms of genes that were determined to be regions specific in a subclass. (**c**) Cell subclasses and genes for which chemotaxis, cell-cell adhesion via plasma membrane adhesion molecules, synapse organization, and axon guidance were enriched terms across the subclasses' region-specific differentially expressed genes. Only genes for which 2 or more cell subclasses were differentially expressed are shown.

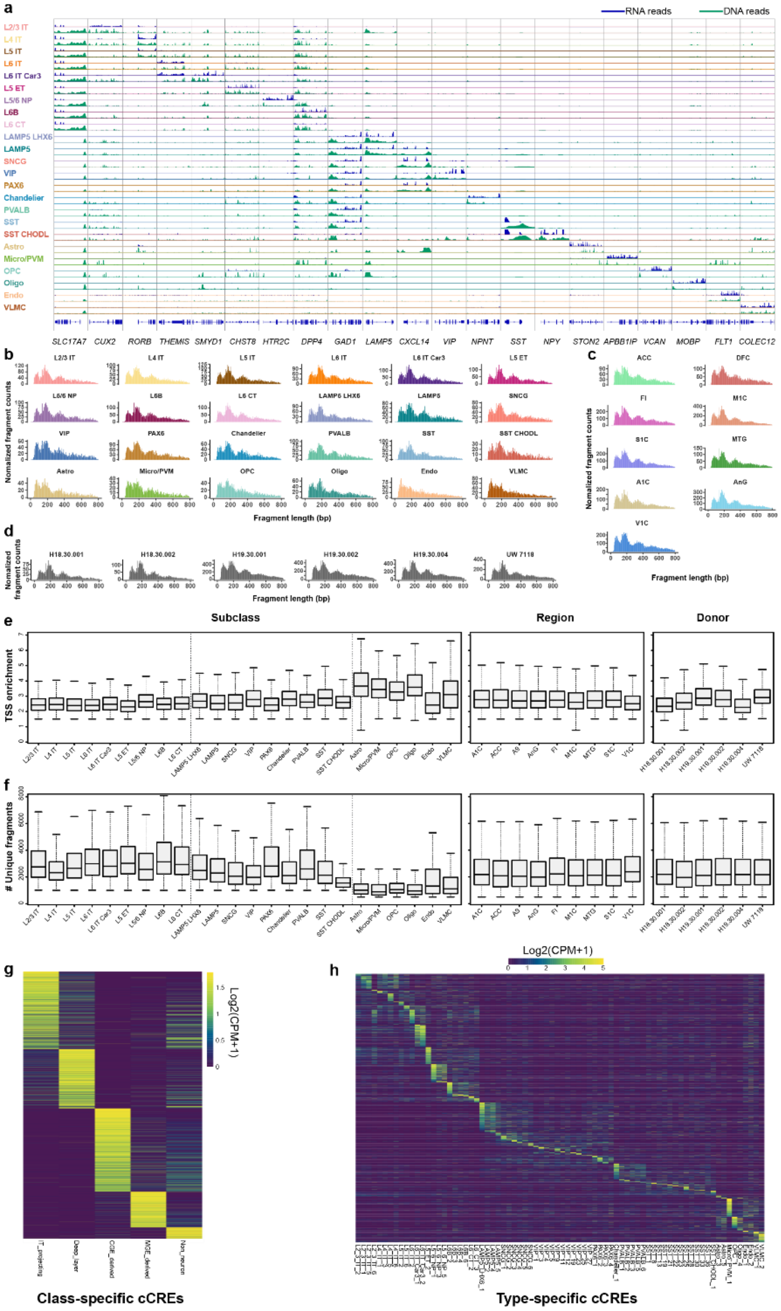


**Supplementary Figure 5. Quality control metrics for single-nucleus ATAC data.** (**a**) Genome browser tracks of aggregate gene expression (blue) and chromatin accessibility (green) profiles for each subclass at selected marker gene loci that were used for annotation. (**b-d**) Fragment size distribution of each subclass (b), each region (c), and each donor (d). (**e-f**) Box plots showing the Transcription start site (TSS) enrichment per cell (e) and number of unique fragments per cell (f) in each subclass, each region and each donor. (**g**) and (**h**) Heatmaps showing the chromatin accessibility of major class-specific candidate cis regulatory elements (cCREs) (g) and cell type-specific cCREs (h).

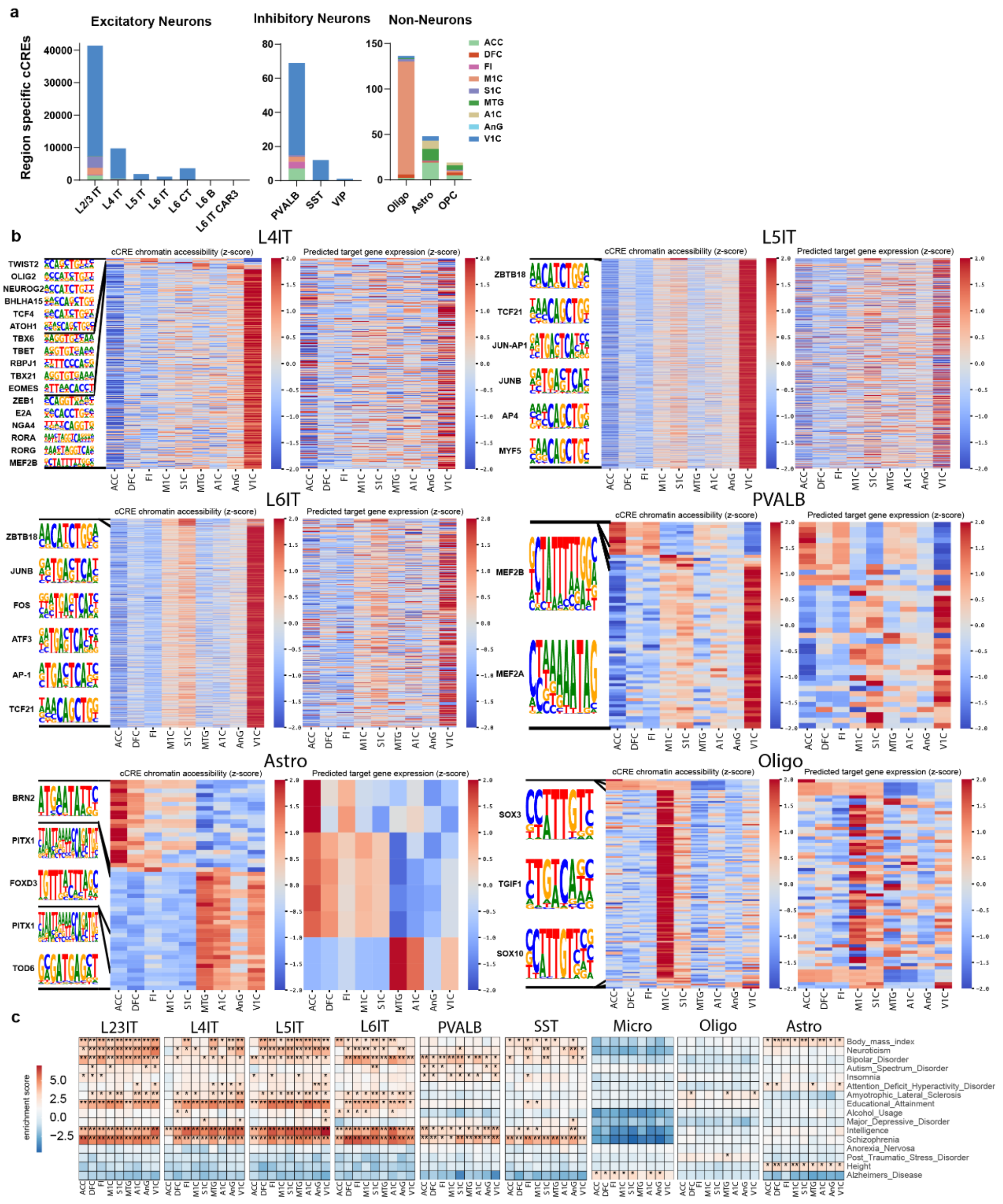


**Supplementary Figure 6. Regional enrichment of accessible chromatin across various cell subclasses.** (**a**) Total counts of region-specific cCREs calculated as chromatin differentially accessible in one region as compared to all others combined. Variable cCREs calculated from all cells that passed filtering. Plots are organized by subclass and colored by region. (**b**) Analysis of regionally enriched accessible chromatin by subclass. Top enriched *de novo* motifs for various subclass region-specific cCREs (left) Heatmap showing the chromatin accessibility of subclass regionally enriched cCREs (center) and heatmap of gene expression linked to subclass regionally enriched putative enhancers as determined by scGLUE (right). (**c**) Heatmap showing enrichment of risk variants associated with various traits from LDSC analysis in subclass enriched cCREs.

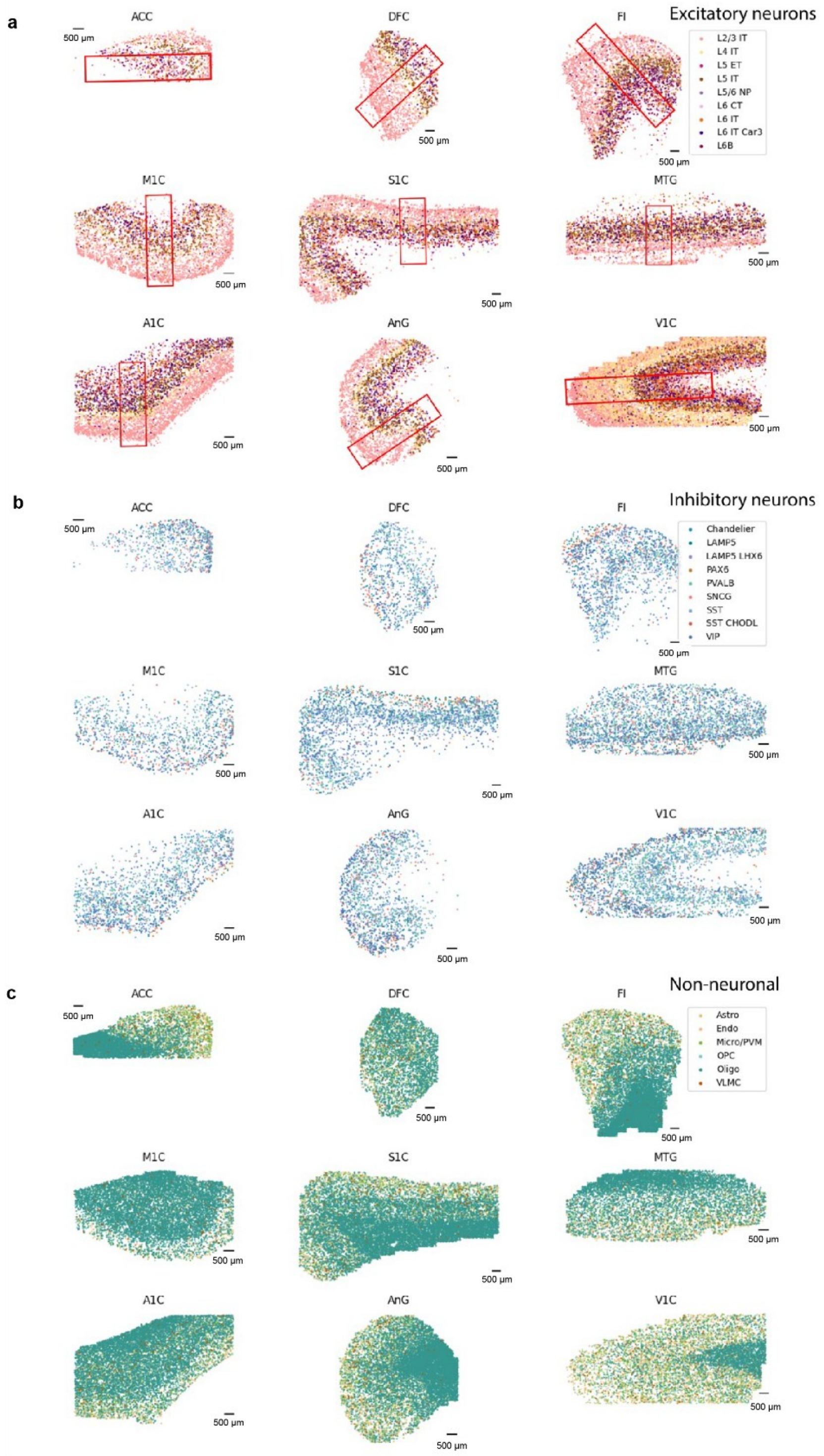


**Supplementary Figure 7. Tissue-wide spatial cellular distributions.** (**a-c**) Spatial cellular distribution across entire tissue sections for excitatory neurons (a), inhibitory neurons (b), and non-neurons (c). Scale bars are 500 µm.

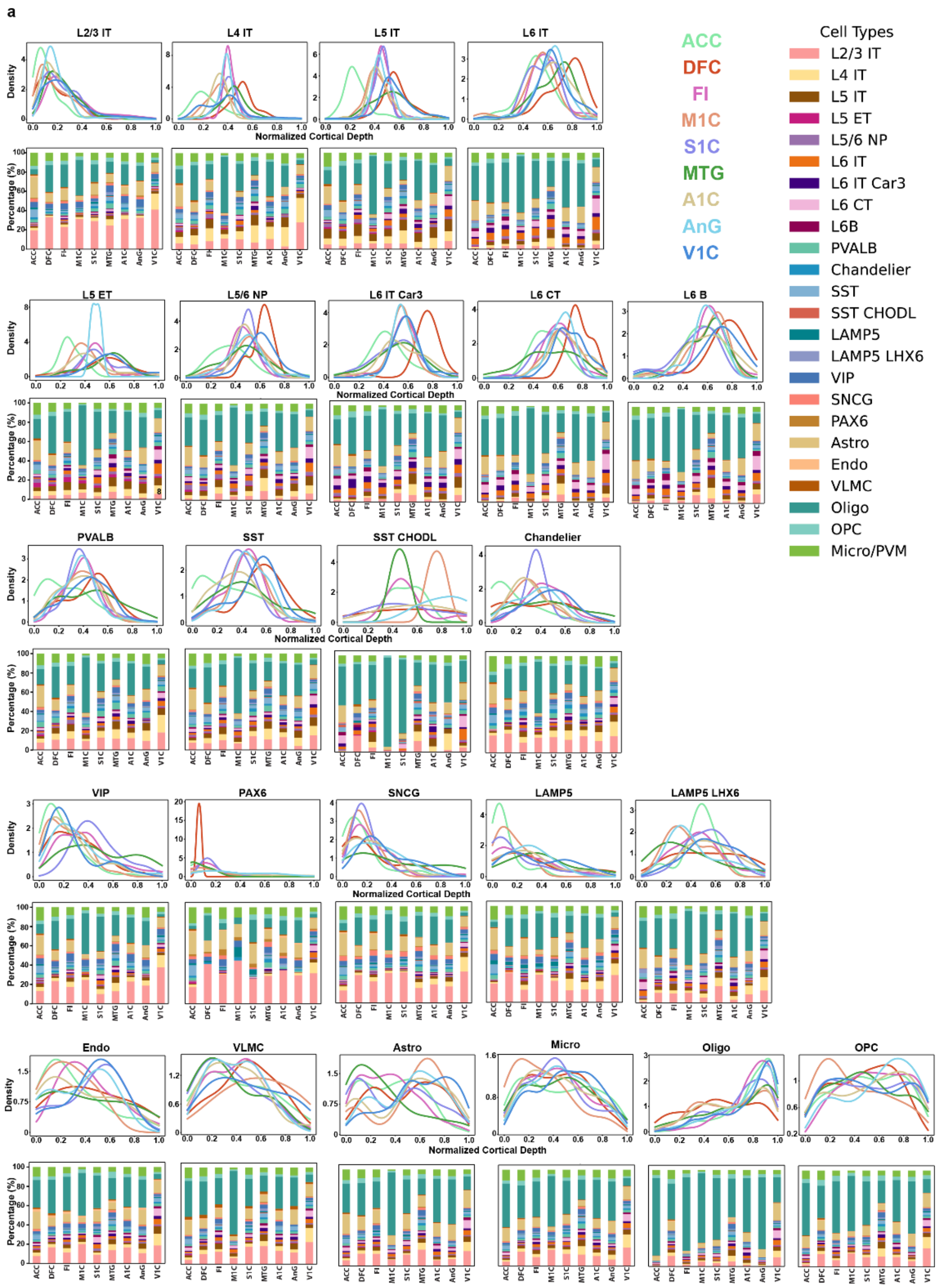


**Supplementary Figure 8. Cortical depth and neighborhood analysis for cellular subclasses** (**a**) Subclass density across normalized cortical depth. For each section, markers of pial surface and white matter were used to establish the cortical sheet and the density of each cell subclass was plotted as a density value across each region. Stacked bar plots visualizing neighborhood analysis for a specific cell subclass. Plotted values represent the percentage of called cells within 200µm of the primary cell subclass averaged across 100 profiled target subclass cells.

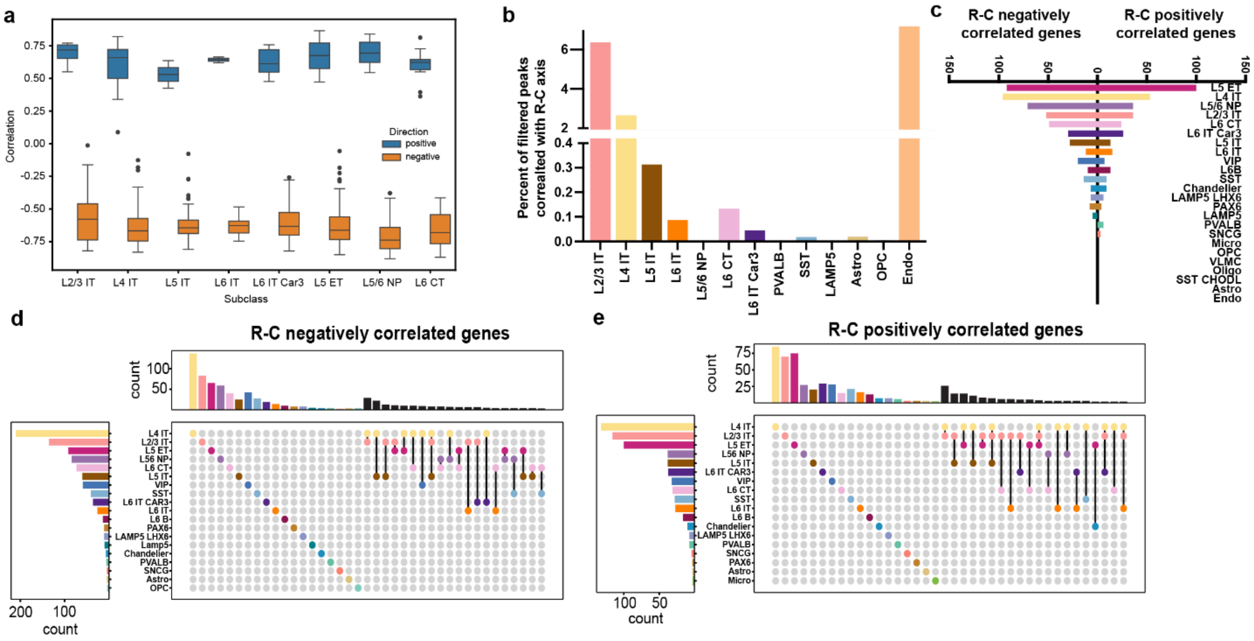


**Supplementary Figure 9. Genes and cCREs correlated across R-C axis.** (**a**) Pearson correlation of genes across the R-C axis from data published in Jorstad et al. Genes selected for the plot are positively (blue) and negatively (orange) R-C correlated genes from this dataset comparisons are completed across numerous subclasses. (**b**) Percent of filtered peaks correlated with R-C axis. The total peaks correlated with the R-C axis as determined in Fig. 4C were divided by the total number of chromatin peaks called in the corresponding subclass across the entire dataset. (**c**) Counts of total genes correlated with the rostral-caudal axis in a uniformly downsampled subset (1000 nuclei/subclass). Correlation is defined as Pearson correlation >0.7 and adjusted *p*-value (BH Corrected) < 0.01. (**d**) UpSet plot of R-C negatively correlated genes. Correlation is defined as Pearson correlation >0.7 and p value <0.01. (**e**) UpSet plot of R-C positively correlated genes. Correlation is defined as Pearson correlation >0.7 and p value <0.01.

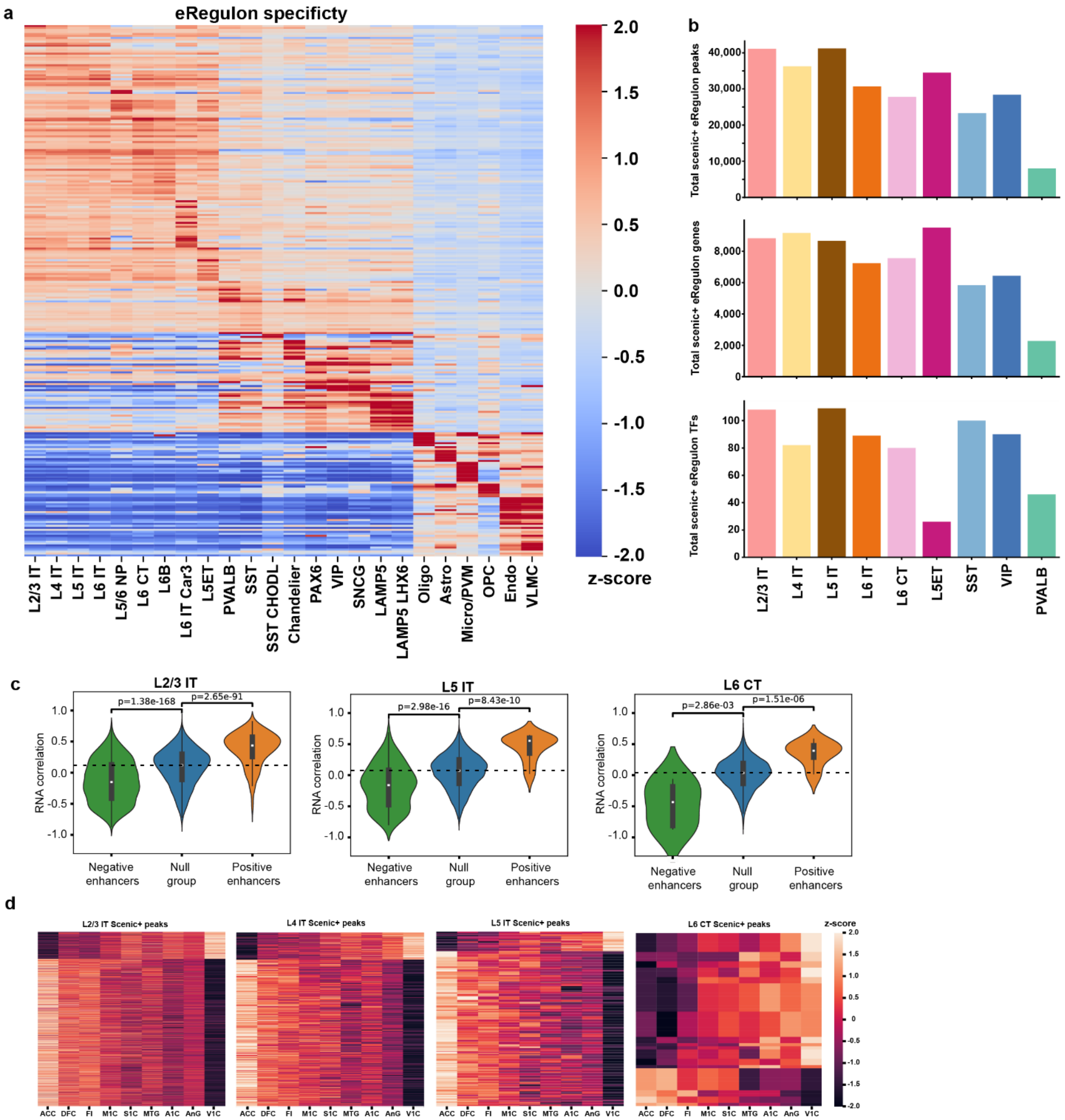


**Supplementary Figure 10. Subclass-specific eRegulons** (**a**) Specificity scores of predicted enhancers that have significant correlation with the R-C axis and regulate predicted target genes in eRegulons. (**b**) Total number of peaks, genes, and transcription factors identified in eRegulons for each subclass profiled via SCENIC+. (**c**) Transcriptomic correlation of predicted target genes from predicted enhancers identified as eRegulon components in neuronal subtypes. Enhancers were classified as positive (orange), null (blue), or negative (green), according to their R-C axis Pearson correlation. p-value from Mann-Whitney U Test. (**d**) Heatmaps of chromatin accessibility by region. Z-scored chromatin accessibility values are plotted for the enhancers predicted by SCENIC+ to regulate genes that have R-C correlated expression.

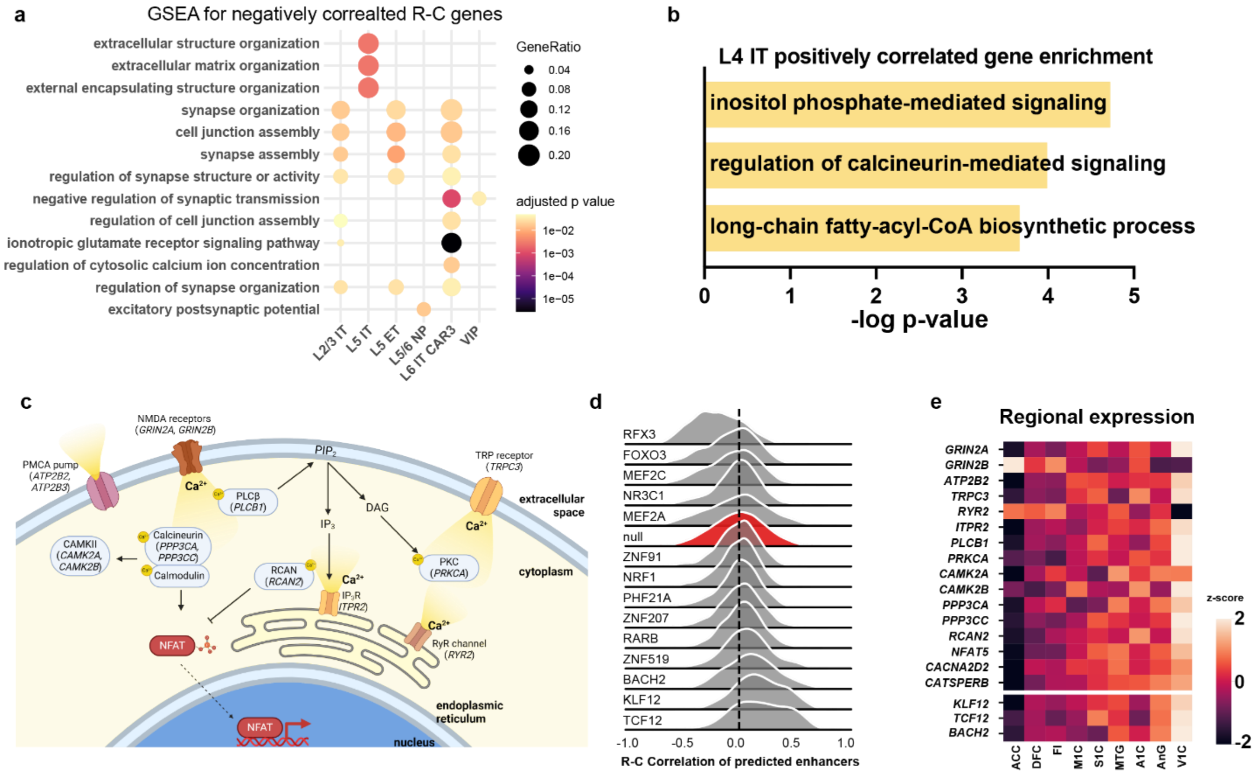


**Supplementary Figure 11. Gene regulatory networks controlling calcium gene expression across the R-C axis.** (**a**) Dot plot showing both gene ratio and adjusted p-value for top enriched terms of genes that had negative R-C correlation in a specific subclass. (**b**) Terms from gene set enrichment analysis of L4 IT positively correlated genes. (**c**) Schematic outlining cellular functions of calcium regulating genes correlated across the R-C axis. (**d**) Distribution of the R-C Pearson correlation of chromatin accessibility of predicted enhancers for transcription factors determined by SCENIC+ to regulate calcium genes shown in Fig. 4H. (**e**) Gene expression of transcription factors and calcium genes by region across R-C axis. Z-score values are shown.

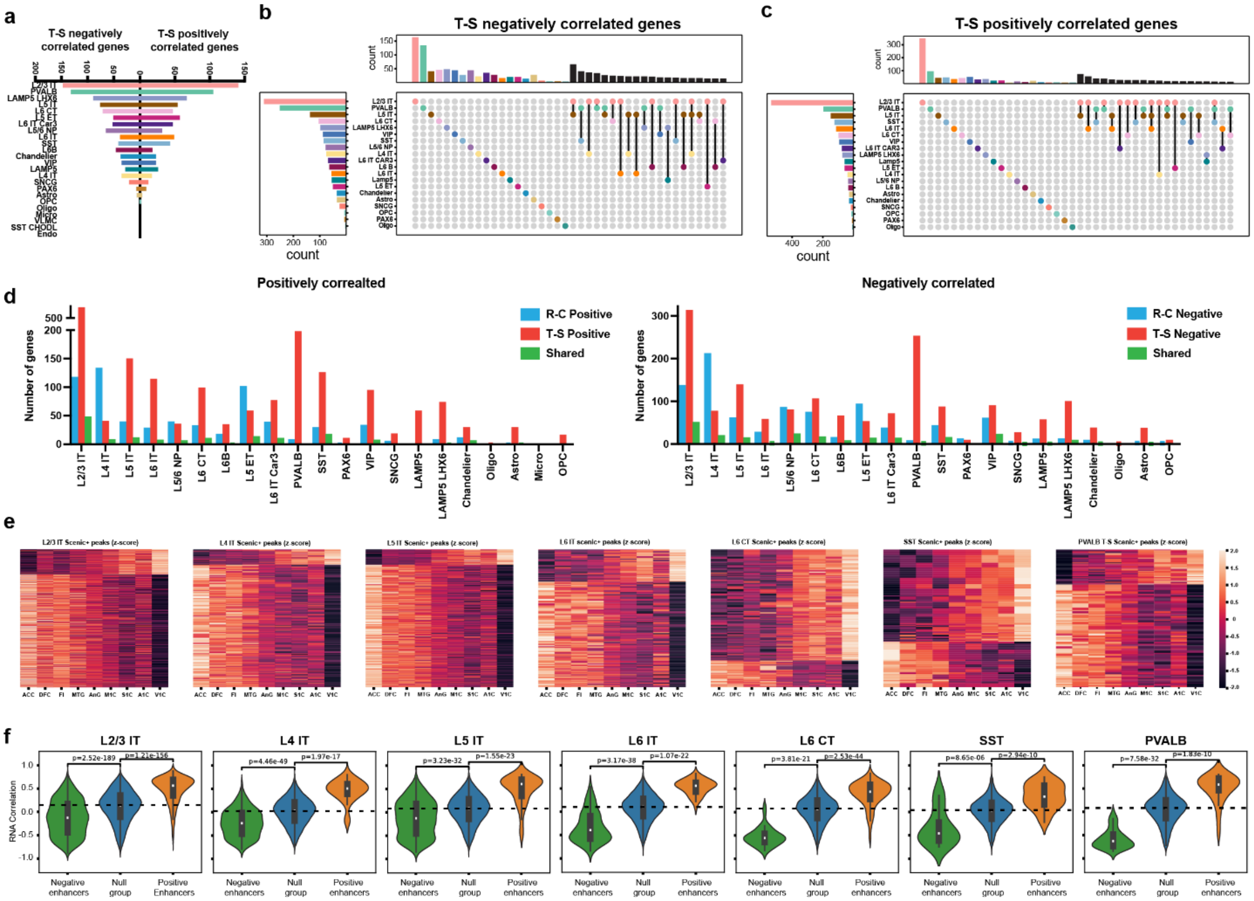


**Supplementary Figure 12. Genes and cCREs correlated across T-S axis.** (**a**) Counts of total genes correlated with the T-S axis in a uniformly downsampled subset (1000 nuclei/subclass). Correlation is defined as Pearson correlation >0.7 and adjusted $p$-value (BH Corrected) < 0.01. (**b**) UpSet plot of T-S negatively correlated genes. (**c**) UpSet plot of T-S positively correlated genes. (**d**) Conservation of genes positively or negatively correlated across the R-C and T-S axis. R-C (blue) and T-S (red) bars represent the total number of genes correlated across that axis while shared (green) bars represent the number of genes correlated in the same direction across both axes. (**e**) Heatmaps of chromatin accessibility by region. Z-scored chromatin accessibility values are plotted for the enhancers predicted by SCENIC+ to regulate genes that have T-S correlated expression. (**f**) Transcriptomic correlation of predicted target genes from predicted enhancers identified as eRegulon components in neuronal subtypes. Enhancers were classified as positive (orange), null (blue), or negative (green), according to their R-C axis Pearson correlation. p-value from Mann-Whitney U Test.

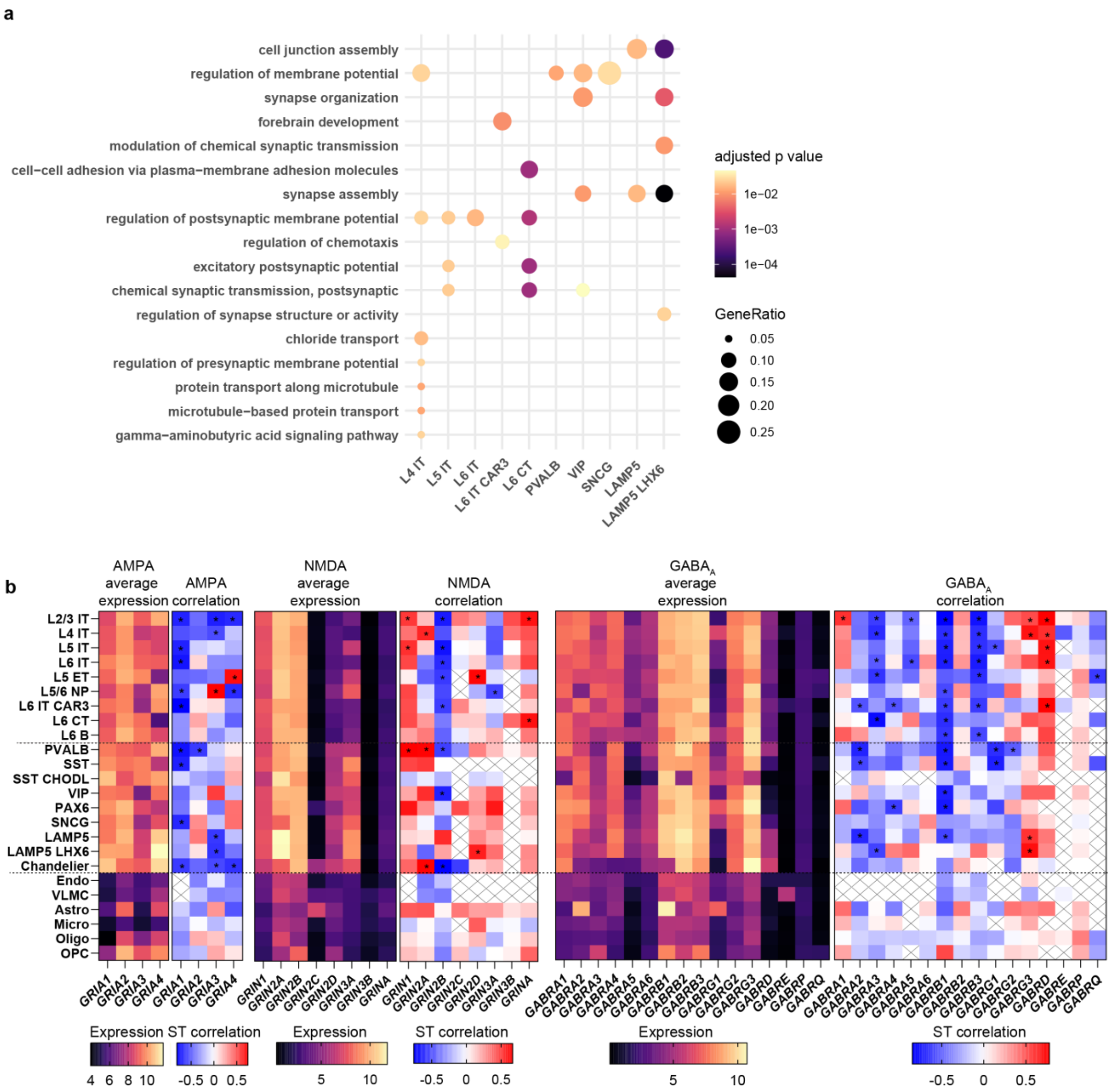


**Supplementary Figure 13. Subunit switching of AMPA, NMDA, and GABA receptor subunits across the T-S axis.** (**a**) Dot plot showing both gene ratio and adjusted p-value for top enriched terms of genes that had negative T-S correlation in a specific subclass. (**b**) Expression values (left) and T-S correlation (right) for each receptor subunit across subclasses. Statistical significance of adjusted p-value of Pearson correlation denoted *p>0.01.

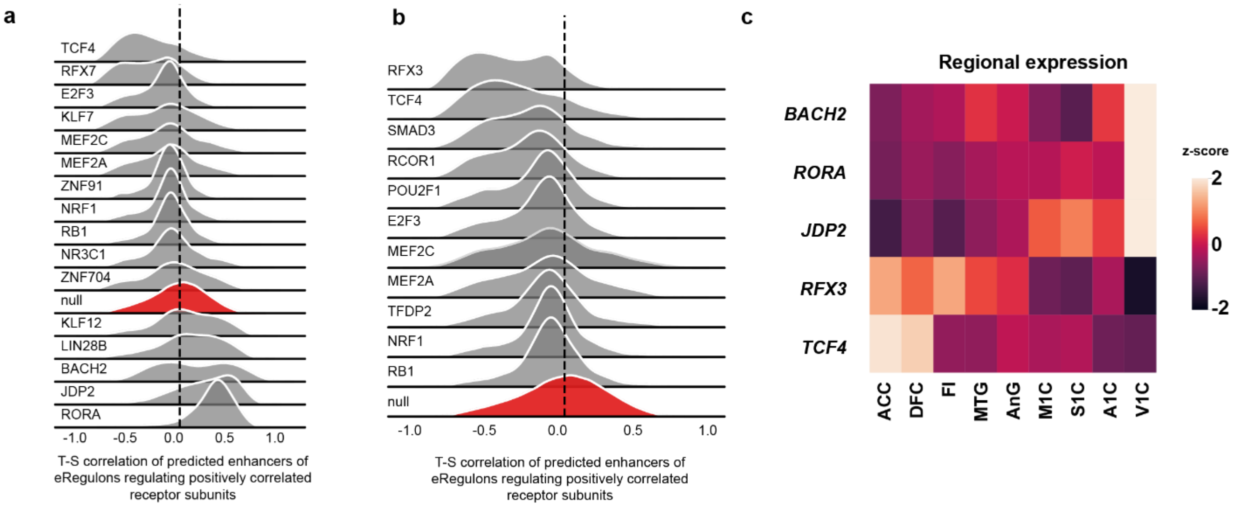


**Supplementary Figure 14. Gene Regulatory Networks controlling receptor subunit switching across T-S axis.** (**a**) and (**b**) Distribution of T-S Pearson correlations of predicted enhancers for transcription factors predicted by SCENIC+ to regulate positively correlated (a) and negatively correlated (b) receptor subunit genes shown in Fig. 5H. (**c**) Gene expression of transcription factors by region across T-S axis. Z-score values are shown.

**Supplementary Data 1. (separate file)**
Donor and sample metadata, subclass information, and subclass marker information by region

**Supplementary Data 2. (separate file)**
List of region specific differentially expressed genes across each cell subtype

**Supplementary Data 3. (separate file)**
List of region specific cCREs across each cell subtype

**Supplementary Data 4. (separate file)**
Panel genes and target sequences for DART-FISH profiling

**Supplementary Data 5. (separate file)**
Correlation of genes across the RC axis by cell subclass

**Supplementary Data 6. (separate file)**
SCENIC+ eRegulon analysis for all cells with subclass specificity scores for each transcription factor.

**Supplementary Data 7. (separate file)**
SCENIC+ eRegulon analysis for select neuronal subclasses

**Supplementary Data 8. (separate file)**
RC activity correlation of identified transcription factors

**Supplementary Data 9. (separate file)**
Correlation of genes across the TS axis by cell subclass

**Supplementary Data 10. (separate file)**
TS activity correlation of identified transcription factors
